# Analysis of spatial and temporal risk of Peste des Petits Ruminants Virus (PPRV) outbreaks in endemic settings: A scoping review

**DOI:** 10.1101/2024.06.21.599995

**Authors:** Julius Mwanandota, Jean Hakizimana, Eunice Machuka, Daniel Mdetele, Edward Okoth, George Omondi, Augustino Chengula, Sharadhuli Kimera, Emmanuel Muunda, Gerald Misizo

**Affiliations:** Tanzania Veterinary Laboratory Agency, P.O. Box 9254, Dar es Salaam, Tanzania; SACIDS Africa Centre of Excellence for Infectious Diseases, SACIDS Foundation for One Health, Sokoine University of Agriculture (SUA), P.O. Box 3297 Chuo Kikuu, Morogoro, Tanzania; International Livestock Research Institute, P.O. Box 30709, Nairobi 00100, Kenya; Ministry of Livestock and Fisheries, P.O Box 2870, Dodoma, Tanzania; College of Veterinary Medicine, University of Minnesota, 1365 Gortner Avenue St. Paul, MN 55108 United States; College of Veterinary Medicine and Biomedical Sciences, SUA, P.O. Box 3019, Chuo Kikuu, Morogoro, Tanzania

**Keywords:** Peste des petit ruminant (PPR) epidemics, spatial methods, spatiotemporal methods, risk factors

## Abstract

**Background:** Sustained Peste des petits ruminants (PPR) circulation, as evidenced by surveillance, shows PPR endemicity in Africa and Asia. Regional transmission of PPR is enabled by joining numerous epidemiological factors. Spatial, spatiotemporal and transmission dynamics analytical methods have been used to explore the risk of PPR transmission. The dearth of information on the risk factors associated with spatiotemporal distribution and transmission dynamics of PPR at a regional scale is high. Through a thorough analysis of peer-reviewed literature, this study sought to evaluate the risks of Peste des Petit ruminant virus (PPRV) epidemics by noting distinctions of geographical and spatial-temporal approaches applied in endemic settings.

**Methods:** A scoping literature review of PPR research publications that used spatial and spatiotemporal approaches to assess PPR risks in endemic areas was carried out using PubMed and Google Scholar data base.

**Results:** Out of 42 papers selected 19 focused on Asia, 15 on Africa, and 8 had a global view. 61.9% used clustering analysis while 35.7% used spatial autocorrelation. Temporal trends were described by most studies at about 71.2% while modeling approaches were used by 13 articles (30%). Five risk factors evaluated include demographics and livestock–wildlife interactions (n = 20), spatial accessibility (n = 19), trade and commerce (n = 17), environment and ecology (n = 12), and socioeconomic aspects (n=9). Transmission dynamics of PPR was covered in almost all articles except 2 articles but it has linked all the risk factors.

**Conclusions:** The review has contributed to the shifting and improvement of our understanding on PPR outbreaks in endemic settings and support evidence-based decision-making to mitigate the impact of the virus on small ruminant populations. Linkage of other risk factors to livestock trade which is the major driver of livestock movement has been shown to pose a significant risk of PPR epidemics in endemic settings. With many studies being found in Asia compared to Africa, future development of predictive models to evaluate possible eradication strategies at national and regional levels should also consider Africa.

## 1. Introduction

A virus known as peste des petits ruminants (PPR) affects small ruminants, primarily sheep and goats, but it can also infect domestic animals. The PPR virus (PPRV) is a single-strand, non-segmented RNA virus of the genus Morbillivirus in the family *Paramyxoviridae* (1). PPRV’s genome spans 15,948 nucleotides (nt) and is structured into six open reading frames (ORFs). The six structural proteins that are encoded by these ORFs are the polymerase(P) or large protein(L), fusion protein (F), phosphoprotein (P), matrix protein (M), hemagglutinin protein (H), and nucleoprotein (N). Additionally, the non-structural proteins C and V are encoded by the ORF transcription unit (2). Four lineages have been described from two structural proteins N or F by phylogenetic studies using partial gene sequences (3) These lineages of PPRV are distributed in several geographical area including Africa, Asia and Europe (4). All four PPRV lineages are present in Africa, where lineage I viruses have been confined in West African countries since 1940. Current evidence shows that lineage I viruses are no longer circulating, as this lineage has not been identified since 2001(5). Lineage II is predominantly present in West Africa, although it has recently been reported in the Democratic Republic of the Congo (DRC) and Tanzania (6). Neither the north nor the west of the continent have reported seeing Lineage III, although reports of it can be found in the Comoros islands and in northeastern, eastern, and central Africa. Africa’s most common lineage, lineage IV, has been documented in fifteen different countries.(6). To date, it has been identified in the northern, western, central and eastern regions of Africa and is gradually moving southwards. Hundreds of millions of domestic small ruminants and wildlife are at risk of infection as the PPRV continues to spread across previously uninfected regions. But the PPRV infection that has been found in previously uninfected areas and the admixture of lineages in countries that have been infected jointly emphasize the geographically and temporally dynamic character of PPR (7).With annual global economic losses estimated at approximately $1.45 and 2.1 billion USD, half of these losses impact Africa, and a quarter affect Asia. These losses are caused by mortality, which reaches up to 20% in naive population and morbidity, which reaches up to 100% (8,9). A global program to eradicate PPR by 2030 has been formally launched by the Food and Agriculture Organization (FAO) and the World Organization for Animal Health (WOAH, formerly known as OIE), due to the high impact PPR on sheep and goat farmers (10).

In the wake of global PPR eradication campaign, several models based on spatial and spatiotemporal risk analysis of PPR epidemics have been adopted to support the eradication programs. Given the existing PPR risk posed by livestock contacts, studies on spatiotemporal models for PPR risk factors and disease transmission dynamics are important for livestock disease prevention (11,12). These models include those related to network analysis, prediction and simulation models. Engaging spatial analysis and identifying areas with PPR clusters, followed by characterising the drivers of the dynamics in those clusters, could generate critical information for disease investigation (13). Enormous availability of such information could support the development of locally adapted control and surveillance strategies (14). Combining such risk factor information with genetic and mobility network data may make it possible to identify disease hot spots, which are crucial for virus entry and dissemination to various regions. (15). Identification of PPR hotspots will involve use of spatial and Spatiotemporal tools to analyze PPR epidemics data. Such analysis will also explore the patterns and risk factors of the disease (16).

Nevertheless, a variety of spatiotemporal tools are accessible for evaluating a range of spatiotemporal hypotheses to meet distinct goals. Either in testing a hypothesis, the outcomes of such a study can be directly compared if the analytical methods used are well understood (17,18). For instance, studies examining how local elements like topography, socioeconomics, demography, and environment can impact disease reporting, identification, and circulation changes over time and space (19). The important step in choosing a model’s parameters is to consider the disease pathway that is thought to connect epidemiological factors with epidemics. (20). In this aspect, disease transmission dynamics is put into context to a specific location where contact network is factored in disease diffusion to various areas (21). The need for this review remains critical because of the overall knowledge gap on effectiveness of these tools in the analysis of PPR control in endemic situations (19).

To our knowledge, no studies have reviewed the spatial and spatio-temporal methodologies used in PPR research. A narrative review by (name the references then cite them) (22) and (23) summarizes the occurrence and distribution of PPR in Tanzania, a systematic review paper by (4,24,25) estimated the prevalence of PPR in sheep and goats and evaluates the potential factors that contribute to the variability in the prevalence and distribution of disease. The only Scoping review conducted by (26) was on PPR diagnostic platforms. The models, variables, spatial, and spatiotemporal methodological frameworks used to model PPR risks were not discussed in those prior reviews. A dearth of information on risks-based spatial and spatial-temporal analysis of PPR epidemics in endemic settings is disreputably very high, and it became scarcer when it comes to infections with multiple hosts (sheep and goat) and high mutation rates (RNA) (27). Filling the aforementioned gap will assist in developing control strategies for PPR, leading to its eradication.

This study aimed to explore the risks of Peste des petit ruminant (PPR) epidemics by evaluating the differences in spatial and spatial-temporal methodological approaches employed in PPR studies in endemic settings. The information obtained from this synthesis process will give an insight into priority research areas and offer suggestions for potential interventions. Moreover, the findings from this review could inform the standardization of modelling inputs and strategies for improved comparability between their outputs.

## 2. Materials and methods

### 2.1. Protocol

A scoping literature review was conducted according to Preferred Reporting Items for Systematic Reviews and Meta-Analyses extension for Scoping Reviews (PRISMA-ScR) guidelines (**Error! Reference source not found.**), in which a checklist for scoping reviews was adopted (28). This study followed the methodology proposed by Arksey and O’Malley (29), identifying the research question, identifying relevant references, selecting references, charting the data, collating, summarizing and reporting the results.

### 2.2. Research questions Identification

The following research questions were the focus of the scoping review.:

a. Which methods, including spatial autocorrelation, have been used to study the spatial and spatiotemporal distribution of PPRV in endemic settings?
b. What is known from the literature in relation to PPR risk based on the analysis of these events (e.g., identification of hotspots, clusters, seasonal or temporal trends)?
c. Which risk factors have been examined and which have been linked to the occurrence of PPR in endemic countries (Africa and Asia)?
d. What is the effect of examined risk factors on transmission dynamics of PPRV?

### 2.3. Search strategy and selection criteria

On the basis of spatial and spatial-temporal analysis, a scoping review of the PPRV risks was carried out. The search strategy included a set of keywords on spatial and spatiotemporal methods, and risk factors in endemic areas were identified with the help of a library specialist for electronic bibliographic search. Peer-reviewed original articles published in English language journals from January 1993 to June 2023 were obtained from systematic searches of two electronic bibliographic databases. MEDLINE (PubMed) and Google Scholar search engines were used as sources of the peer-reviewed articles included in this review. The articles were selected using keywords combined by Boolean operators’ peste des petit ruminant virus (PPRV) AND risks AND spatial and spatial-temporal analysis AND epidemics AND endemic countries. While searching PubMed, key keywords were reduced to generate results. The searches from the two electronic databases hit a total of 766 records (Google Scholar: 648, and PubMed 118 as in Figure 1.

**Figure 1:**
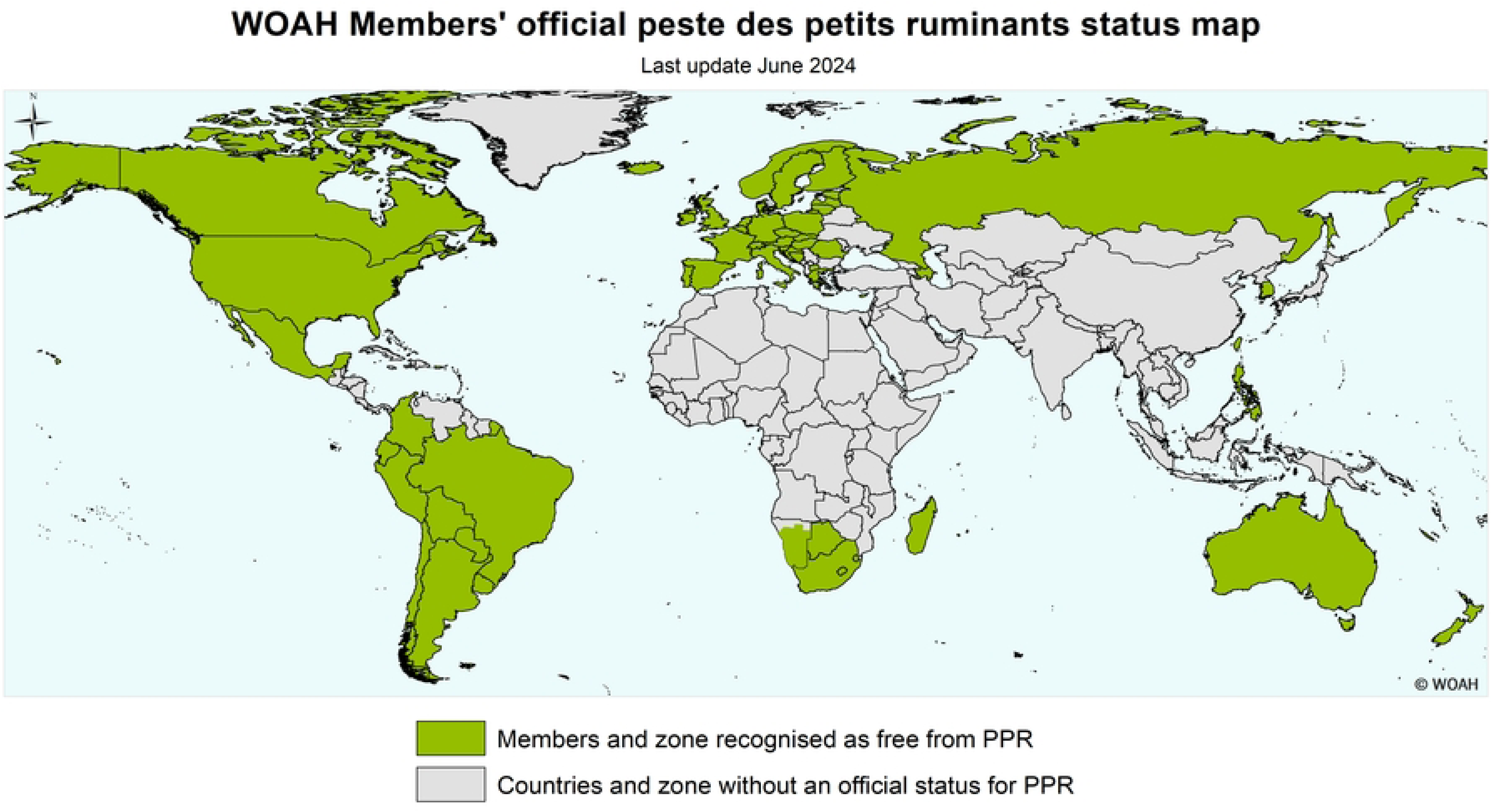
Peste des petit ruminant (PPR) situation based on World Organization for Animal Health (WOAH) PPR status recognition. Regions in gray are endemic, report sporadic episodes or have an unregistered PPR status.

### 2.4. Eligibility criteria

This scoping review included PPR studies incorporating spatial, temporal or spatiotemporal approaches for data analysis and inference. Based on the classification proposed by (18) studies, different Spatiotemporal analytical tools can be used for epidemiological research. These can be classified based on the purpose of the analysis on (a) visualization and description, (b) spatial or Spatiotemporal dependence and pattern recognition, (c) spatial smoothing and interpolation and (d) modelling and regression studies. Within the category of modelling and regression, we were particularly interested in studies designed to investigate epidemiological factors linked to PPR outbreaks at the population level or that integrated presumed PPR risk factors to project disease occurrence. Inclusion criteria were done by selecting all papers with original peer-reviewed articles. The PPR endemicity was defined based on the information available at the latest WOAH list of members and zones recognized as free from PPR (Resolution No. 17 (89^th^ General Session, May 2022 (30); Countries recognized as PPR free with or without vaccination as shown in Figure 1 were excluded from the analysis. However, studies conducted in countries certified as PPR-free or with recognized free zones were eligible for inclusion, for example, Russia (31). Thus, studies considered for inclusion were those that (a) reported PPR epidemics distribution/patterns or modelled PPR risk using geographically or/and temporally linked data; (b) used population outbreak data collected in PPRV endemic countries; (c) were published in a peer-reviewed scientific journal between January 1990 and July 2023 in English and (d) documented PPR occurrence in livestock (e.g., cattle, small ruminants, camels) and wildlife.

### 2.5. Exclusion criteria

Studies that used data collected from farm outbreak surveys with self-reported PPR occurrence and risk factors were excluded. Also, studies that were designed as outbreak investigations, risk assessments and narrative literature reviews documenting PPR occurrence and trends (without further data analysis) were out of the scope of this synthesis. Studies from countries recognized as PPR-free with or without vaccination were excluded from the analysis.

### 2.6. Selection of relevant and reliable studies

A search strategy was developed and run in three electronic bibliographic databases and truncated free-text terms were combined using Boolean and proximity operators to obtain a good balance between sensitivity and specificity while searching for the studies. Electronic search outputs were imported to Mendeley. The reports were first screened using the titles and abstracts, and duplicate studies were eliminated. At this stage, irrelevant papers were excluded. Full texts were retrieved to assess the final inclusion based on the eligibility criteria. An individual reviewer read the full texts before deciding which ones to include. For the purposes of traceability and transparency, the reasons for exclusion following this stage were recorded. By applying the eligibility criteria, two reviewers with backgrounds in PPR surveillance and molecular epidemiology screened the articles for selection. The first selection was from a title and abstract screening, and the second one was from a full-text screening. All conflicts generated through the screening stages between the two reviewers were discussed until a consensus was reached.

### 2.7. Data extraction from included studies

Every study’s data was extracted by a single reviewer using a pre-made, standardized form in a Microsoft Excel Spreadsheet that contained the following:

a. study characteristics – author, year of publication, years analyzed, country, geographical coverage, species.
b. PPR outbreak data and diagnostics – diagnostic criteria, data source, and surveillance system;
c. the type of analytical tool used, the process for grouping and identifying patterns in the data, the method for assessing the data based on time and season, and the list of epidemiological factors that were investigated and their outcomes. One author (JJM) extracted the data, and another (JNH) validated them to ensure accuracy prior to the quality appraisal phase. All included papers were subject to a full-text analysis using a thematic analysis to develop an evidence matrix, which captured relevant data from each source using the main themes emerging from the literature. Themes captured included references to spatial distribution, livestock trade, weather conditions (dry/wet), climate change, and species/age/sex.

### 2.8. Collating, summarizing and reporting the findings

The approaches to formally assess the presence of clusters were classified according to the dimensions and data forms proposed by (17,32). A modified version of the framework proposed by (18) was used as a guide to classify the analytical tools used for spatial or spatial-temporal analysis based on their purpose, as previously described. The factors examined in the epidemiological models commonly represented aspects with the potential to influence PPRV introduction, transmission, survival or features with a presumed role in the effectiveness of control measures. These factors were organized into a framework that included five main categories: (a) spatial accessibility;(b) livestock demographics and livestock–wildlife interactions; (c) livestock trade; (d) socio-economic factors; (e) ecology and environment. Epidemiological covariates that did not fit in any of the categories were grouped separately as ‘other factors’ (**Error! Reference source not found.**) and Figure 4:. With an emphasis on the connection between epidemiological factors and the occurrence of PPR epidemics, all of the data gathered from the studies was presented narratively by combining the key components linked to the review questions, which included study and outbreak characteristics, analytical techniques, and their findings. All analyses were conducted using R packages in R version 4.3.3, which included *ggplot2*, *webr, tidyr* and *dplyr,* to plot figures and to produce quantitative descriptions of the data while maps were developed by QGIS (33–36).

## 3. Results

### 3.1. General characteristics of the selected studies

Following the electronic search approach, a total of 56 studies were included for the qualitative synthesis and 42 for the quantitative synthesis. 101 papers were evaluated in full text from the initial pool of over 766 publications that were screened, and 57 of those were eliminated because they did not meet the eligibility requirements (Figure 2).

**Figure 2:**
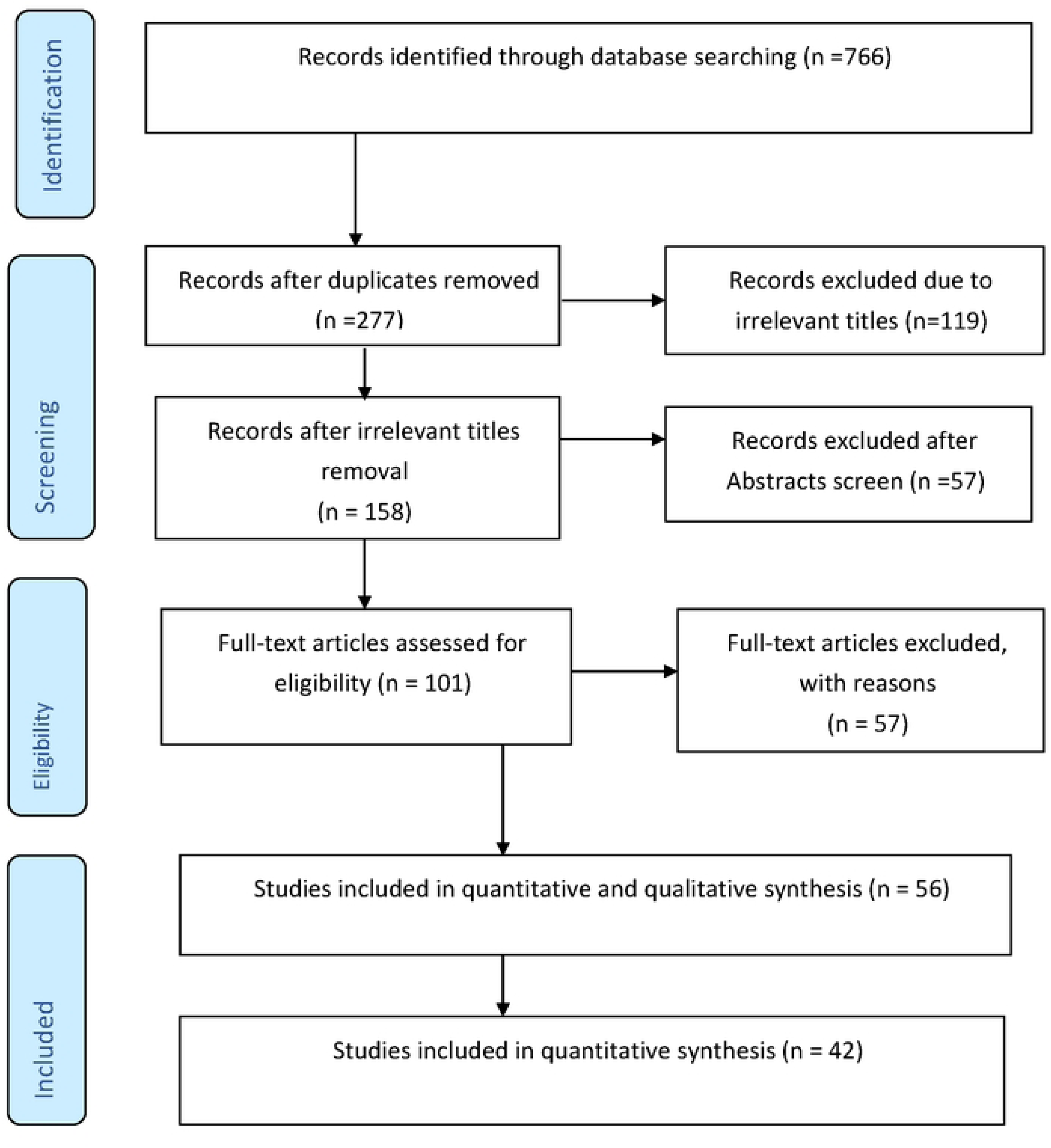
Flow diagram from bibliographic search of study records to final inclusion in the review.

### 3.2. Time intervals and geographic regions

The distribution of studies by geographical scale and scope varied across Africa and Asia (Figure 3). Included studies covering 21 countries distributed across two continents (Asia and Africa). The largest number of studies identified were from China (n = 13), Bangladesh (n = 5) India (n = 4) Ethiopia (n =3), Tanzania (n = 3), as in Figure 3. Most studies were conducted at a national scale using country-wide PPR epidemic data (45%) available through official sources (95%). In most studies, data were collected as part of passive surveillance systems (37.5%) in which PPR epidemics were commonly diagnosed based on clinical presentation only (15%) (**Error! Reference source not found.**). The spatial unit used for analysis and the livestock species for which outbreaks were recorded varied across studies (**Error! Reference source not found.**). Some studies (50%, n = 20) included epidemic reports from sheep and goats. However, other species category analyzed PPR epidemics at the range of 2.5-5% of studies recorded. In some studies, long-term epidemics data was analyzed, the longest being a 44-year PPR epidemics case series from India (37). However, the study period varied across studies (median = 3 years; range: 1–44), with 11 studies analyzing outbreak data collected for more than 10 years. The majority of reports included a mix of analytical tools and objectives, with most studies concentrating on the visualization and description of outbreaks.

**Figure 3:**
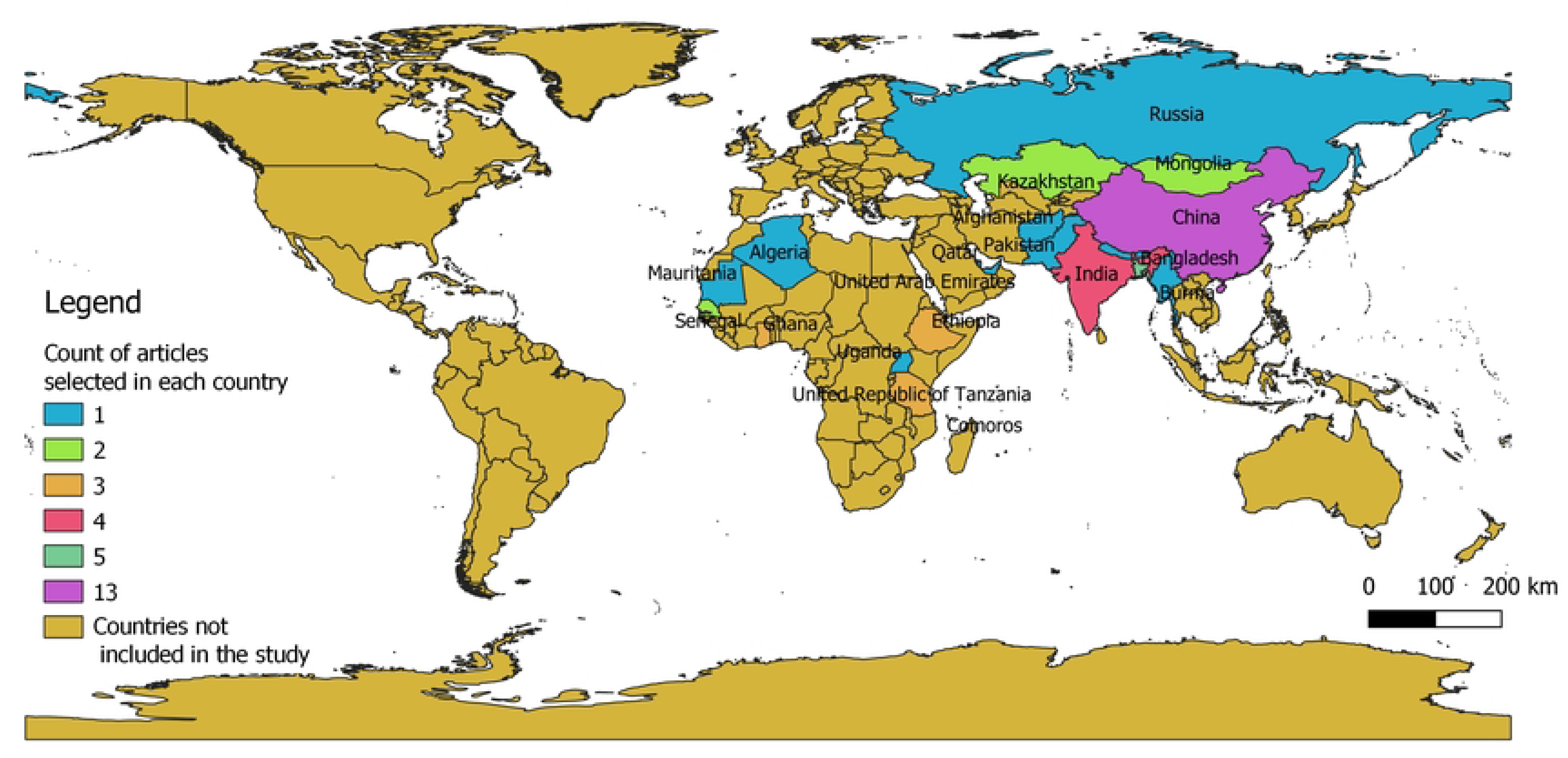
Geographical coverage of the studies included in this review. The map shows the number of studies included per country; countries in Central and West Africa were underrepresented despite being endemic for PPR.

### Methods used to identify spatial and temporal variations of PPRV risk factors

**Error! Reference source not found.** and Figure 5 shows various spatial, temporal and Spatiotemporal methods that were used to visualize patterns, explore spatial clusters, and model risks across space and time. Although the results of some studies suggested the use of these techniques, they did not state clearly how useful they were (38). More than half of the studies (61.9%, n = 26) used at least one spatial clustering analysis technique for testing the non-randomness hypothesis of spatial or spatial-temporal distribution of PPR epidemics; of these, 35.7% (n = 15) and 26.2% (n = 11) used either a spatial or a spatial-temporal tool. Four studies reported the use of various approaches for the identification of spatial or spatial-temporal clusters in parallel. Spatial clusters detected through Moran’s I (n=4), Maximum entropy spatial statistic (n = 5), followed by Getis-Ord (n = 1) and Clark Evans test (n = 1) were frequent, whereas Moran’s I was often applied as a test for areal spatial clustering (n = 4). SaTScan was chosen for simultaneous detection of space-time clusters (n =5) in all studies, whereas temporal and spatial autocorrelations were investigated (11). Spatiotemporal directionality tests to detect the direction of the progression of PPR epidemics were used in three studies (11,39,40).

**Figure 4:**
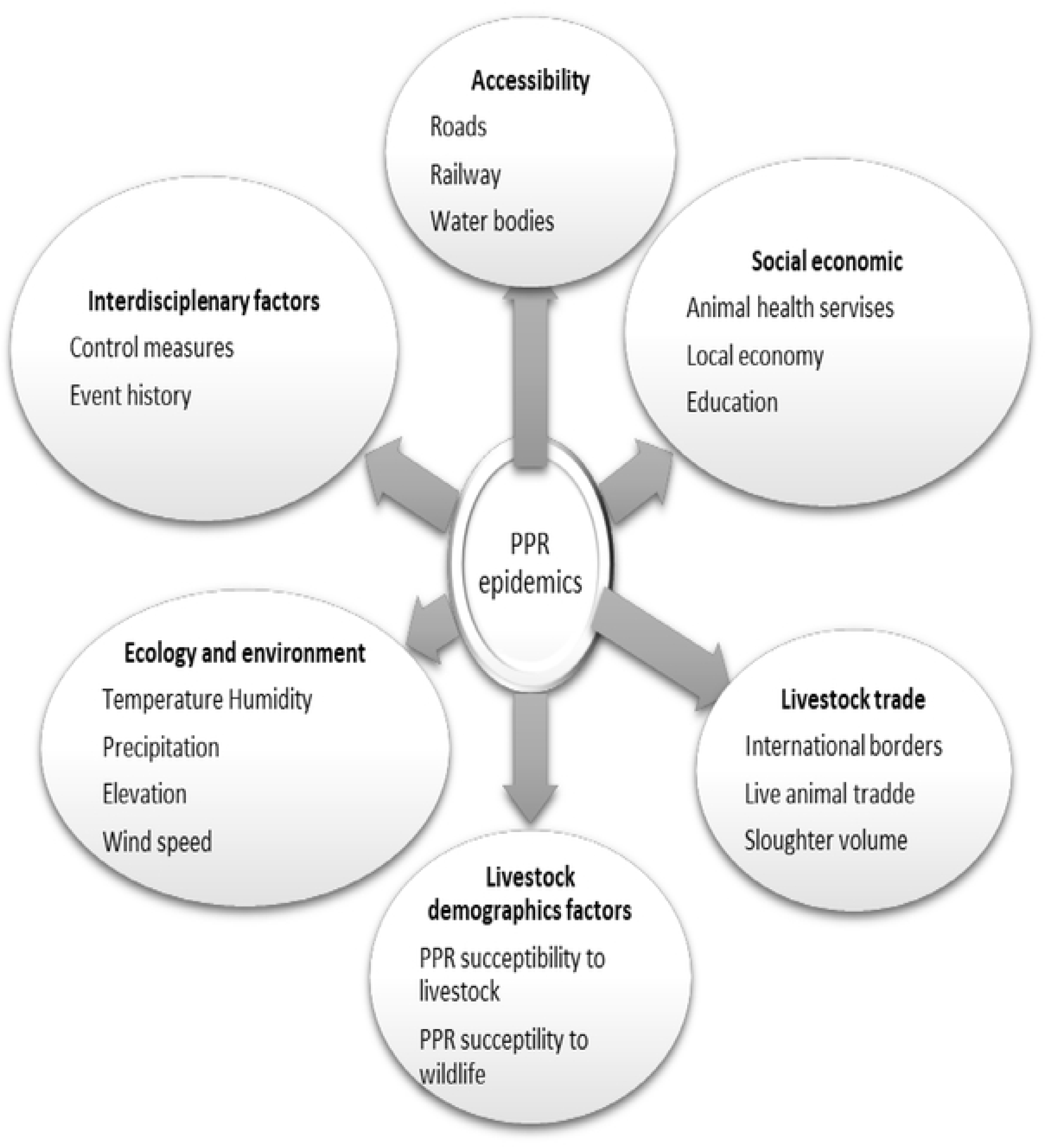
Conceptual frameworks used to classify the risk factors for Peste des petits ruminants epidemics.

**Figure 5.**
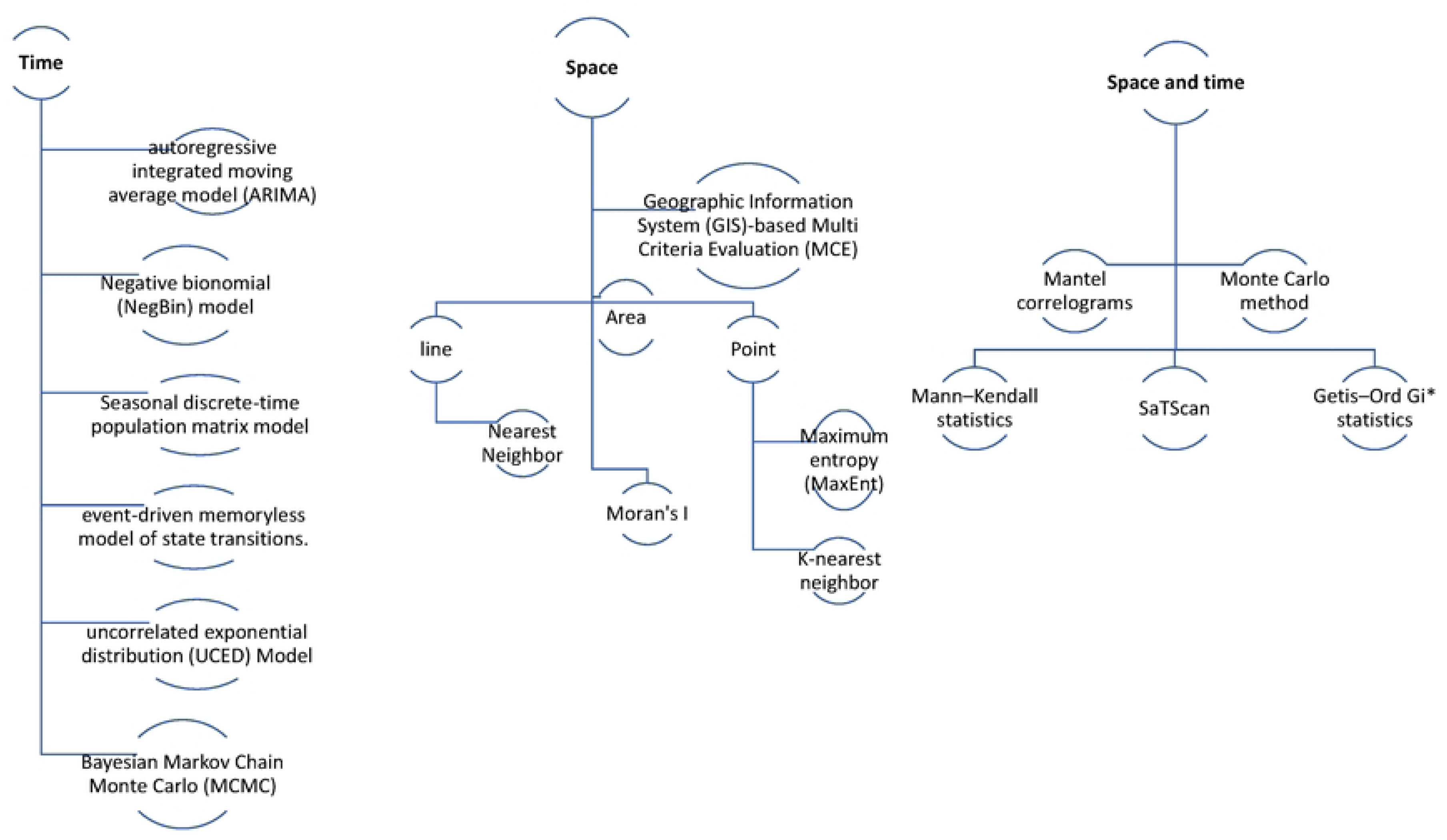
Dimensions and data forms for classification of cluster analysis modified from (41)

### 3.3. Spatial autocorrelation or spatial clustering

Evidence of PPR spatial autocorrelation methods for disease cluster identification was documented in 35.7% (n = 15) of the studies. However, due to the diversity of techniques used to identify such clusters (unusual aggregation of epidemics) and hotspots (excess level of epidemics in comparison to a threshold level), their diverse assumptions and the configuration of the data on these results tended to vary within and across studies(19). Spatial analysis is a useful tool for studying the distribution of infectious diseases, which, due to their transmission dynamics, often follow diverse spatial patterns and commonly occur in spatial clusters(42). Four studies reported heterogeneous results, recording the identification of random or clustered patterns that varied across the periods or analytical methods. For example, (43–45) reported that annual variation influenced the identification of spatial association. The following two studies (11) and (46) detected both clustering tendencies and random spatial patterns, which varied across the years, and the various clustering methods used in the analysis. The methods used for cluster evaluation in the reviewed studies and their general findings are abridged in **Error! Reference source not found.**. Cluster size was measured through the radius of the significant clusters, and their duration varied greatly within and across studies. The latter scenario might reflect both the diverse status of disease at the local level and changes in the tools used, the hypothesis tested and the assumptions adopted.

### 3.4. Temporal and seasonal trends assessment

Most studies (71.2%, n = 30) described the temporal trends of PPR epidemics aggregated per day, week, month or year depending on the temporal aggregation of the original data. Generalized Linear Negative Binomial Regression Model (GLNBRM), (11), Generalized Linear Mixed Models (GLMMs) (47), Negative Binomial (45), linear (15,37,48,49) or logistic regression (50,51) models were used to explore or test hypothesis related to temporal trends. Other studies resorted to Bayesian approaches (3,52,53). The NAADSM (North American Animal Disease Spread Model (54,55), autoregressive integrated moving average model (ARIMA) (56), Least Cost Path (LCP) (57), Random forest (44) Ensemble Algorithm (58), Event-Driven Memoryless model of state transitions. (59), Regression Tree Models (44) and Mantel Correlograms (15), which were employed to investigate the relationship between genetic distances and various measures of network and geographic distance. Moreover, 4 studies formally analyzed PPR seasonality through the assessment of seasonal trends distributed geographically and socioeconomically (31,37,43,60)

### 3.5. Modelling approaches

In addition to describing the patterns of PPR epidemic distribution and pinpointing disease hotspots using clustering techniques, various spatial and spatial-temporal tools were employed to assess the correlation between a number of epidemiological factors and PPR epidemics. Among those studies included in this review, 13 (30.9%) used covariables to explore the link between population-level epidemiological aspects and the epidemics, to predict PPR risk or to produce risk probability maps. 11 studies used epidemiological factors to model or project PPR counts (19%, n = 8) (11,16,39,45,51,53,56–58,61). Six studies focused only on predicting PPR epidemics using epidemiological factors as covariates to forecast the PPRV suitability area and estimate the spatial risk (number of outbreaks in PPR passive surveillance data (40,43–45,58,62). Transmission dynamics were also evaluated using metapopulation model (53). Before applying statistical models to evaluate the risks of the PPR epidemic, visualization and descriptions of numerous PPR epidemics were conducted. These visualizations and descriptions provided a perfect environment for understanding the distribution of the PPR epidemic over space and time. Data visualization and descriptions were used by 34 studies (80.9%), and this was the most reliable method used by all the studies to present the distribution of PPR epidemics (55). Various analytical approaches used for data visualization and description include GLMNB (11), GLMMs (47) and negative binomial to assess the potential risk factors associated with the case count of PPR disease from each outbreak. (45,60). Either linear (15,37,48,49) or logistic regression (50) models were used to explore or test hypotheses related to temporal trends. GeoDa 1.14.0 was used to perform Geographically Weighted Regression (GWR), with climate and geographical variables acting as predictors and log-transformed PPR cumulative incidence serving as the dependent variable.(46). Bayesian techniques, such as Bayesian Time-Scaled Phylogenetic Analysis, were used in other investigations.(3,52,63) along with the empirical Bayesian kriging (EBK) technique for risk map generation and geostatistical prediction (34). NAADSM (North American Animal Disease Spread Model (54), autoregressive integrated moving average model (ARIMA) (56), least cost path (LCP) (57), The Naïve Bayes (NB) and Random forest machine learning algorithms (44), ensemble algorithm(58), regression tree models (44)and Mantel correlograms (15) which was employed to examine the relationship between genetic distances and various measures of geographical and network distance. Seasonal population matrix model was used to assess the ability of different vaccination schedules if they can be incorporated into PPR control program (64). Hot Spot analysis and Cluster analysis were made to identify the hot and cold spot areas by using ArcGIS v10.4.(60). Maximum Entropy Ecological Niche (MaxEnt) modelling was employed to detect suitable areas for PPR virus distribution (40,43). GIS-based multi-criterion decision analysis was used to identify areas at risk of PPR occurrence and spread (62). Event-driven model of PPR developed from susceptible-exposed-infectious-recovered (SEIR) model was used to study how the virus propagates in memoryless state transitions in Afghanistan (59).

### 3.6. Epidemiological risk factors associated to PPR epidemics

Even though the categories of trade and commerce (n = 17), environment and ecology (n = 12), animal demographics and livestock–wildlife interactions (n = 20), spatial accessibility (n = 19), and commerce and trade (n = 17) accounted for a greater proportion of the epidemiological features used in the models, nine studies (n = 9) reported on the role of socio-economic factors on PPR risk. The majority of the models examined the cumulative impact of risk variables that fall into several framework categories (**Error! Reference source not found.** and Figure 6). When there is at least one variable in the group that is either positively or negatively connected with the chance of PPR outbreaks, the result for each risk factor category is labeled as linked. When counting the number of studies conducted in the region, the same number of studies in Africa and Asia examined the impact of various factors in their models, with the exception of socio-economic aspects (Africa: 8/16; Asia: 3/18), animal demographics and livestock-wildlife interactions (Africa: 7/16; Asia: 7/18), livestock trade (Africa: 9/16; Asia: 10/18), spatial accessibility (Africa: 6/16; Asia: 7/18), and environment and ecology (Africa: 3/16; Asia: 4/18). Two studies that were not examined have also been analyzed, out of the six that examine risk variables from a global viewpoint (**Error! Reference source not found.**). Furthermore, epidemiological parameters were incorporated in various forms across studies, indicating a consistent risk pathway linking them to the outbreaks. **Error! Reference source not found.** lists all of the covariables that were examined in detail.

**Figure 6:**
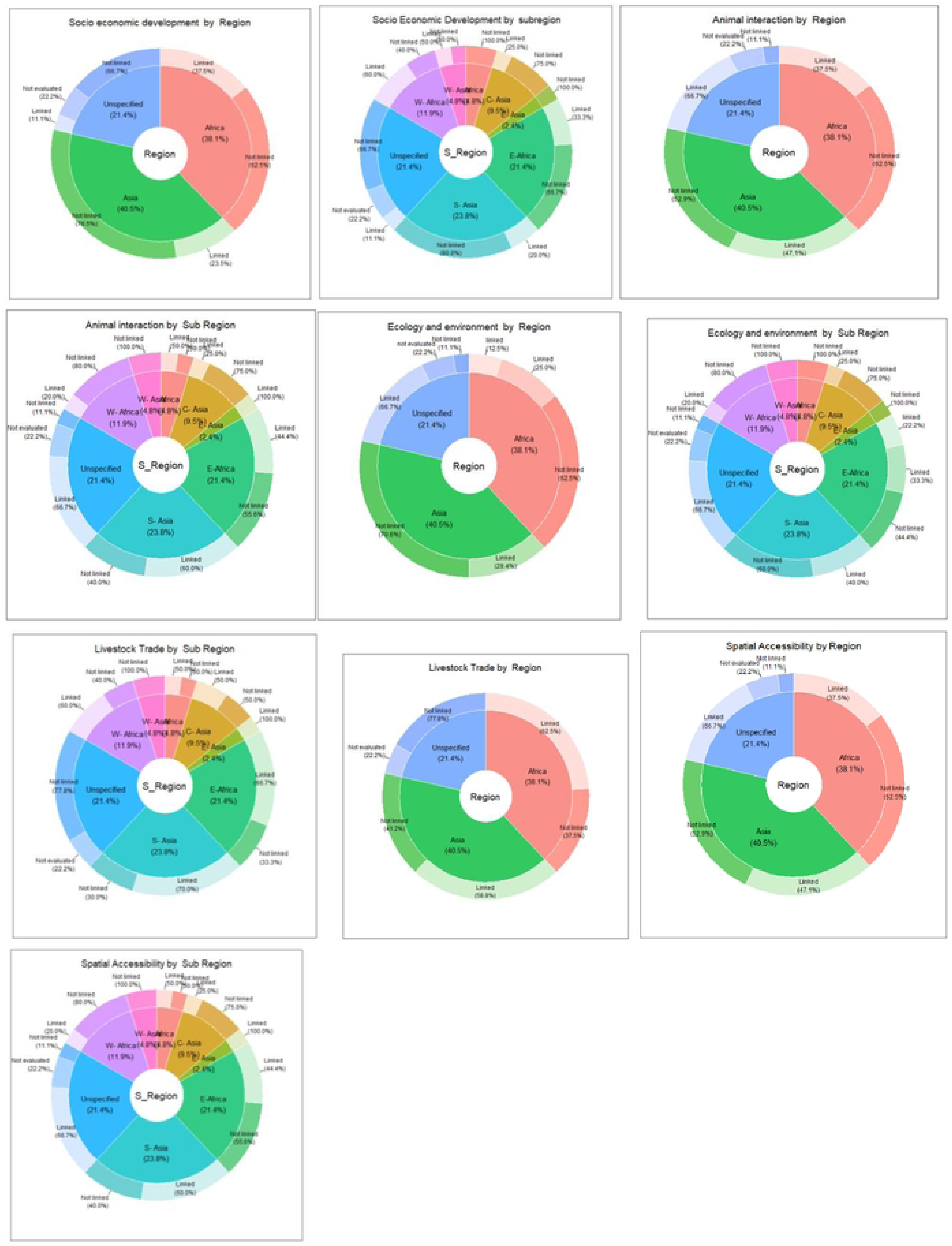
Distribution of models, at regional and subregional, containing variables from every category in the conceptual framework.

#### 3.6.1. Livestock demographics and livestock–wildlife interactions

According to **Error! Reference source not found.** the primary objective of twenty studies (six in Africa, twelve in Asia, and one worldwide) was to identify hotspots by examining the impact of susceptible livestock populations on PPR risk. Sheep and goat populations were reported as a covariate more frequently than other susceptible species, despite the fact that many other species were taken into consideration (e.g. camel, cattle and pigs)(11,16,57,58,62,65–68,27,39,43,45,46,51,53,56). In 3 out of 17 studies, there was a correlation between a small ruminant population size and sheep and goat density, or a higher risk of PPR (27,51,62) whereas the linkage between other species (pigs, camel, wildlife and cattle) population and PPR epidemics varied across studies with more than half (9/17) of the studies reporting an association with the risk of PPR epidemics (9,11,27,43,51,58,62,63,65,69). Merely describing a correlation between wild populations and PPR outbreaks has been the focus of limited research (3/9) (27,63,69,70).

By including covariates that reflect the circulation of PPRV in wildlife reservoirs like saiga goats, an epidemiological connection between PPRV in wildlife and disease outbreaks in livestock was investigated. For example, closeness to protected, national parks or forest coverage were included in seven analyses (four in Africa and three in Asia) as a representation of a potential livestock–wildlife interaction; among these studies, four reported a positive association between presence or closeness to a protected area and PPR outbreak occurrence; however, this relationship varied across study periods, and locations (11,43,47,57,62,63,69). In general, 47.4% (7/19) of research done in Asia and Africa linked this category of wildlife-livestock interaction to PPR risk.

#### 3.6.2. Livestock Trade

Livestock trade-related factors were included in 19 studies, 9 in Africa and 10 in Asia, as in **Error! Reference source not found.**. Probing the effect of international connections on PPR risk through proxy variables that represented the likelihood of an international trade network or cross-border movements was a common objective, with eight studies (three in Africa and five in Asia) incorporating either the distance or adjacency to an international boundary into their models. As reported by six studies, an increased distance to an international border reduced the risk of PPR. (11,15,45,57,69,71). Additionally, elements pertaining to the dynamics of the market were taken into account, such as the movement of live animals or their products. Finding the markets in China, Ethiopia, Uganda, and Comoros was crucial for determining the outbreaks in those countries.(9,11,39,60,62,67). Two investigations examined the relationship between PPR epidemics and human demography (43,71).

#### 3.6.3. Accessibility and networks

Fourteen studies (five in Africa, seven in Asia, and two globally) explored variables that represented landscape and topographical features, including permanent transport networks (e.g., roads and railways) and water bodies (e.g., rivers and lakes) to study PPR risk **Error! Reference source not found.**). All studies evaluating the role of accessibility in African countries (5/5) reported an association between spatial accessibility and risk of PPR; in contrast, six studies (out of seven) analyzing this aspect found an association in Asia. Greater PPR risk was found in locations near major transportation routes or in areas with dense road networks, while shorter distances to roads or railroads were linked to lower PPR occurrence.(11,62). A study from Ethiopia reported that the number of effective contacts per unit of time (watering points or pasture and through live-animal trade) was one of the main predictors for PPR transmission (53). Furthermore, four studies evaluated the impact of natural landscape elements, such as the proximity or extent to inland waterbodies and rivers; only one of these studies noted the buffering effect of adjacent water bodies against PPR occurrence (43).

#### 3.6.4. Association between environmental factors and PPR epidemics

Although 13 studies (3 in Africa, 6 globally and 4 in Asia) assessed the risks of PPR outbreaks due to variations of environmental and ecological features, the review listed 10 features categorized in environmental and ecological factors (**Error! Reference source not found.**). The most frequently analyzed features were season, temperature and precipitation (31,37,43,44,48,58,60,71). Landscape was also assessed by five studies (40,43,49,69,71). Seasonality and landscape were highly linked to PPR epidemics (5/13) in each feature. The diverse results were obtained for weather and other climate-related covariables. Three studies explored the association between elevation and PPR risk, with lower altitudes being highly linked to PPR epidemics compared to higher attitudes (43,45,71). Temperature and precipitation, which determine the status of land cover as humidity, solar radiation and wind speed, have also been shown to influence PPR outbreaks (43,44,58). In general, all of the studies carried out in Asia (4/4) and Africa (3/15) found a correlation between the risk of PPR and ecological and environmental features; however, the variables chosen to investigate the risk differed amongst the studies, illustrating the different analysis goals.

#### 3.6.5. Social and economic development

Socio-economic data was only included in 9 studies (**Error! Reference source not found.**) (50) and (45) to assess the risks of PPR spread in the world and the Republic of Kazakhstan, respectively, based on socio-economic and geographic parameters like landscape indicators. Two studies (43) and (62), one in Asia and the other in Africa, focused on literacy rate and the provision of veterinary services (e.g. the number of veterinarians, technicians and animal health workers). Poor animal health and veterinary services assessed in Mandi City, northwest of India, have increased the risk of PPR infection (43). The contribution of the urban population to PPR epidemics was reported by two studies, one in Africa and the other in Asia (43,47).

### 3.7. PPR transmission dynamics and respective control measures

To illuminate the important PPR transmission dynamics, evaluation of PPR risks was done in almost all studies except two (9,17), which have not explicitly evaluated this covariate. Four studies have explicitly evaluated the transmission dynamics in relation to the control (51,53,56,61). The first one was carried out in Ethiopia and estimates the level of PPRV transmission in endemic settings using a metapopulation model that simulates PPRV spread.(53). The second one uses a mathematical model to assess the impact of four vaccination strategies implemented at different times in reducing the PPR burden (61). In India, Susceptible-Exposed-Infectious-Removed (SEIR) were used to evaluate PPR transmission dynamics (56). Another study conducted in Tanzania has provided evidence of the PPR transmission pattern (51). However, the probability of PPR transmission is linked to some covariables evaluated in this review; only 13 studies are influenced by livestock contact. Livestock movement as the main contact parameter in PPRV transmission has been evaluated using network analysis to identify PPRV hot spots(15). Those covariables included temporal and spatial aspects, production systems and the use of PPR control strategies such as vaccination and early reporting systems (8) (43). Moreover, all six studies evaluating the impact of vaccination reported a decreased risk of PPR outbreaks in areas participating in PPR vaccination programs (9,53,59,61,64,72). Despite the indisputable significance of PPR transmission dynamics, most studies have used temporal and spatial resolution during epidemiological data collection. The diverse range of these linked covariables has included all epidemiological parameters in this review.

## 4. Discussion

The epidemics of PPR have substantially impacted the economies of many African and Asian countries. Spatial and spatiotemporal approaches have played a critical role in shifting and improving our understanding of the available disease management options. However, the uneven load may be attributed to the diverse data sources, covariates and analytical approaches employed. The results of our analysis show an increasing trend in the number of studies applying spatial and spatial temporal analysis for estimating PPR risks in endemic countries. The most frequent analytical tool employed in this review was those related to data visualization and descriptive exploratory analysis using georeferenced data. Descriptive results are the foundation of the epidemic analytics pipeline; the hypotheses generated at this stage normally precede and guide the development of more complex analyses and models that seek to gain in-depth knowledge on the aspects influencing temporal or spatial disease variation (18,73).. The progression from descriptive to analytical was a key feature of several study reports included in this review. Assessment of PPR transmission dynamics has improved our understanding on the means of disease risks tracking, control planning and eventual elimination of the PPRV.

### 4.1. Modelling approaches

Methods deployed in analysis of spatiotemporal distribution in this review has used both tools for visualization and descriptive analysis. These tools have shown the extent of spatial distribution by means of size, shape, and directionality of PPR spread. Through the later information our understanding on extent of PPR impact in endemic countries has been improved significantly. The knowledge of these methods can assist in the designing of PPR control strategies which include assigning vaccine or surveillance buffer zones and recognizing the distance closest to risk factors. Based on descriptive analytical tools evaluated in this review, planning of PPR control strategies can be easily supported. As more data becomes available, model development and refinement remain vital for understanding risks of PPR epidemics in local settings. Most importantly, predictions of PPR epidemics need to be updated to account for livestock mobility patterns on the fundamental spread of PPR in order to reflect the heterogeneous risk in space and time.

### 4.2. Spatial autocorrelation

Spatial autocorrelation or spatial dependence, is a key component of PPR spatial epidemiology (43). Quantification of spatial autocorrelation in some studies was done by using global spatial autocorrelation indices (18,74) i.e., Moran’s I (43) Mantel test (15), and Getis Ord(45). Importance of analyzing spatial temporal autocorrelation is critical for seasonal and temporal trends disease evaluation. Either outbreak, cluster and hotspot detection can offer hints to the concealed causes of the increased incidence of diseases and associated drivers for its endemicity(17). The usefulness of these results is not limited to hotspot detection but it goes further to PPR targeted control measures (proactive or reactive vaccination, culling or depopulating and/or quarantine).

### 4.3. Temporal trends

Epidemics of PPR like other diseases tend to have seasonal variation due to continuous changes of environmental and ecological factors. In our review we have identified how these environmental and ecological factors underlying seasonal transmission of PPR are critical for predicting and understanding the long-term environmental trends effects of livestock health (75). PPR spread is influenced by animal contacts, resource sharing, economic activities, and trade. Seasonal variations in precipitation, temperature, and availability of pasture and water can affect livestock mobility and PPR risk projection. Seasonal activities like festivals and dowries during harvest also increase the risk of PPR spread. Control strategies include strategic vaccination and movement restrictions before disease outbreaks. These risk factors vary geographically and demographically.

### 4.4. Risk factors associated with PPR epidemics

In this review several categories of risk factors which explore demographics (human and livestock) and livestock contacts have been evaluated in many studies. Source, type and extent of interaction of animals has been the major focus of evaluation. Variability of sheep and goat density in each category has influenced the magnitude of each risk category. Livestock movement which account for most of PPR outbreak in highly accessible area has been substantiated in this review. Livestock trade which is the major driver of livestock movement is associated with increased risk of PPR epidemics in highly accessible area. Geographical and environmental predisposition of sheep and goat production may influence the population of small ruminant in any given area. Increased density of wildlife or livestock in the interface area may increase the risk of PPR spread between livestock and wildlife population. Wildlife livestock interaction can increase the number of PPR epidemics if the population of PPR host animals is significantly high. Socioeconomic activities which provoke interaction of people and animals account for PPR risks in area with high population of sheep and goat. Identifying PPR-prone areas is crucial for putting risk-based surveillance and control measures in place.(62).

#### 4.4.1. Livestock-wildlife Interactions

The impact of livestock–wildlife interactions on the risk of PPR epidemics is linked to the density of susceptible species at livestock wildlife interface area. In our review several studies have shown evidence PPR outbreak in a protected area to be due to its close proximity with PPR risk area. Few evidence of PPR in wildlife species in Africa (76) is surprising given the increasingly visible epidemics in wildlife in Asia (69). Evidence from our models suggest that the spatiotemporal patterns of PPRV outbreaks in wildlife were similar to those in livestock, suggesting evidence of PPR virus spillover from livestock to wildlife (77). Global climate changes have resulted in shifts in species distributions and habitat suitability consequently reduced resource availability and increase wildlife–livestock interactions. In our models evidence of increased risk of PPR infection due interfered relationship between species, habitat and climate leading to mortalities of wildlife species e.g saiga (*Saiga tartarica tartarica*) in Kazakhstan has been demonstrated (70).

#### 4.4.2. Accessibility and network

Livestock movement has been the main focus in this risk factor because it is the main driver of contacts between livestock population. The majority of the models evaluated have demonstrated that PPR outbreaks are connected with geographical links provided by infrastructure.(62). Large Water bodies like lakes, river and ocean has also been evaluated to identify evidence PPR risk. High accessibility to these features facilitates contacts of animals and human through trade and other socioeconomic activities. Poor infrastructure and proximity to water bodies as seen in mountainous area and island respectively may limit spread of diseases. Expansion of vehicular long-distance transportation of live animals, limited accessibility to pasture and water resources and the enroute insecurity of non-vehicular transportation may have reduced enroute disease contamination but enlarge the radius of disease spread.(78). By combining genetic and mobility network data, our review has clarified models that assess the risk of PPR transmission resulting from livestock movement and identify critical sites for virus entrance and spread in particular regions. The analysis is critical to improving our ability to use risk factors and data on livestock movement to create locally tailored control and surveillance strategies.(15)

#### 4.4.3. Social economic Factors

The impact of socioeconomic factors on the risks of PPR spread in this review has been evaluated in diverse models with focus on political, economic, veterinary services and stakeholders’ knowledge. Lack of scientific data on the spread of diseases due to the political situation in some nations, such as Afghanistan, has led to unsuccessful large-scale mitigation strategies. A herd-level, event driven PPR model was constructed to study the viral dissemination among a flock. A model like this can identify effective management strategies for different herd compositions and circumstances. The model has given decision-makers a tool to create efficient PPR epidemic containment measures. To offer an update on the significance and impact of PPR economically, a cost-benefit analysis adopting various control techniques was assessed. Some models assess the accessibility of veterinary services in relation to PPR hazards, which were also correlated with the national economic status. Stakeholders’ knowledge assessment has shown to be critically useful in involving livestock keepers in planning and implementation of control and eradication program. Furthermore, sheep and goat are kept by vulnerable population (women, Childrens displaced population). Engagement of vulnerable population like women through increased access to vaccines and other control packages will support PPR eradication programs (79). Social economic vulnerability was not limited to gender inequality as for woman above but there were other factors like Climate change. Resilience to climate change among pastoral and agropastoral society were the drivers of PPR spread. Risk mitigation strategy of PPR include livestock mobility, herd diversification, livestock selling, increasing drought livestock, and shifting to another place. In our analysis, we were able to demonstrate the relationship between PPR outbreaks and socioeconomic elements that encourage host interaction, including the trade in livestock, cultural events, husbandry methods, nomadism, and economic and ecological factors. (80,81). Other social practices such as livestock marriage dowries can also promote the spread of PPR to other areas through the livestock mobility(80). Social and political status of PPR endemic country may influence future regional transmission dynamics hence should be highly considered during plan for control and eradication. In cost benefit analysis quantification PPR control benefits is critical for resource mobilization and political will towards PPR eradication.

#### 4.4.4. Livestock trade

The growth of Sheep and goat production has been associated with increase in transhumance and consequently transboundary spread of infectious diseases such as PPR. In this review we evaluated market-related dynamics of sheep and goat linked to PPR risks. The evaluation was done by looking the likelihood of livestock trade to facilitate spread of PPR in the respective area. Livestock trade infrastructure like road and railways network, slaughter facilities were examined for evidence of PPR risks. For instance, in Bangladesh, the livestock movement within herds and the central market of the local small and large markets has been connected to the PPR outbreak (46). Another model which evaluates urbanization and habitat characteristics has shown naked evidence to PPR risk to livestock trade(47). Urban areas are endowed with different level of improved Livestock trade infrastructure (transportation network and slaughter facilities) varied in different regions. In this review the identified regional variation in trade related PPR risks call for different approaches while designing control strategies in local context. International Market dynamics were linked to PPR spread where outbreaks in the middle east, Europe and China were traced back to sub-Saharan countries. Empowering sub-Saharan countries with livestock trade infrastructure which support PPR surveillance and control is critical for meeting 2030 PPR eradication target.

#### 4.4.5. Ecology and environment

Studies on landscape and climatic features linked to PPRV environmental circulation, stability, and survival has been assessed in this review. Results has shown geographical variation in risk magnitude. For example, area with good precipitation will have good pasture and water supply (40). Shrinkage of Grazing land due to factors like expansion of conserved area, agricultural activities and climate change has been linked PPR risk. In this review we have been able to identify prediction models which can be used in designing control strategies according to the environment and ecology of respective area. For example in China seasonality pattern identification is critical for vaccination schedule because during summer high temperature lower animal immunity (57). The ecological niche model discussed in this study has demonstrated how ecological and environmental characteristics are related to PPR outbreaks. The annual maximum temperature was inversely correlated with PPR outbreaks because PPRV is more susceptible to hot temperatures. Other elements, such as seasonal variations in precipitation (warm vs. dry season), exhibited a substantial positive connection with PPR outbreaks, whereas wind speed had a negative association.(58). Prediction models allow us to identify the right time for control measures deployment, it become ineffective if done outside the time range hence 2030 PPR eradication target has been identified using prediction models(58,82).

#### 4.4.6. PPR transmission Dynamics

The dynamics of PPR transmission depend on the rate of transmission from PPRV infected animal to susceptible hosts. In disease models, this rate is of transmission is captured in one parameter called probability of transmission (83). In our review several studies have captured spatial transmission dynamic modeling approaches to investigate PPR transmission dynamics and control (56,84). These models have been useful in generating scenario analyses of the potential course and severity of PPR epidemics by characterizing and forecasting the spatiotemporal transmission patterns of PPR epidemics, or assessing the effectiveness of interventions and the feasibility of achieving elimination targets(37). Our review have been able to integrate key epidemiological features of PPR infection and strive to capture relevant mechanisms of PPR transmission, including the potential influence of environmental factors (85). In metapopulation models, a particular type of spatial dynamic model where the population is divided in a set of interacting population groups defined according to spatial or demographic information (53,86). In Ethiopia for example, PPRV incursions into highlands from lowlands are due to uneven distribution of livestock markets, pasture, and water, requiring vaccination targeting interfaces between different populations locations (53). Our understanding in PPR transmission dynamics have the potential to achieve high level of communication speed between the response team in the outbreak event. Using the knowledge on PPR transmission dynamic the outbreak will evoke a coordinated events from the point of origin and contributors, instantaneous on-the-ground epidemiologic characterization, evaluation to establish spread pathways, specimens to molecular characterization of the virus, while performing spatial prediction models (86).

### 4.5. Strength

Most likely, this scoping review is the first to offer details on risk-based spatiotemporal techniques connected to PPR spread in endemic situations. The results of this research indicate a promising trend in the scientific communities’ use of spatial epidemiology tools to comprehend the PPR’s spatial transmission mechanism. While spatial and temporal analysis has been extensively used, our evaluation indicates that the quality of these studies and the analytical methods vary by investigation. Our evaluation gives a roadmap of the work that has already been done in the field and identifies areas for future research towards improving and creating spatial approaches for investigating PPR. This evaluation, in contrast to the preceding review study by (4,23,24,26) also gains from the inclusion of a number of methodological restrictions of existing spatial studies that may limit the ability to provide reliable data to support local control initiatives aimed at lessening the burden of PPR. Lastly, we feel our assessment gave a fair depiction of PPR risk mapping initiatives since we used a thorough search method in compliance with the PRISMA Scoping criteria.

### 4.6. Limitations

We acknowledge the challenge of modeling PPR risks in endemic situations given the lack of real data for model parameterization and validation and the inherent complexity of the employed approaches. Some models are intentionally hypothetical, where the focus may be more on appropriate mathematical tools, in an effort to provide new ideas or offer a fresh perspective on a critical process for PPR transmission and spread. Prediction models based on metrological data and simulation models based on current data sets are two examples.

It should be highlighted that even though the data extraction and screening were carried out separately by two independent reviewers, it is possible that we overlooked certain publications that were relevant to our study’s objectives and may have made significant contributions. Because only studies published in English was included in our review, pertinent papers published in other languages may have also been overlooked.

### 4.7. Conclusion

Spatial and Spatiotemporal approaches have played a critical role in shifting and improving our understanding of the available disease management options for PPR. However, the uneven burden of PPR may be attributed to the diverse data sources, covariates, and analytical approaches employed. Future development of prediction models to assess potential eradication efforts at national and regional levels should also take Africa into consideration, as many studies have been conducted in Asia as opposed to Africa.

## Declarations

### Ethics approval and consent to participate

Not applicable

### Consent for publication

Not applicable

### Availability of data and materials

The datasets used and/or analyzed during the current study are available from the corresponding author on reasonable request.

### Competing interests

The authors declare that they have no competing interests

### Funding

Through the Regional Scholarship and Innovation Fund (RSIF), which Julius Joseph Mwanandota was given to use for his doctoral studies at the SACIDS Africa Centre of Excellence for Infectious Diseases, SACIDS Foundation for One Health, Sokoine University of Agriculture, Morogoro, Tanzania, the Partnership for Skills in Applied Sciences, Engineering and Technology (PASET) funded this study.

### Authors’ contributions

- **Julius Joseph Mwanandota (JJM)** formulated the research question, carried out the literature search, and synthesized the literature, all of which contributed to the study’s design, data collecting, and interpretation of the results. Write the article first, then give it a critical edit. He assumed the major position in the Manuscript’s writing as well.
- **Jean Hakizimana (JNH)** evaluated and edited drafts of the written report, helped with the synthesis of the literature and the interpretation of the study findings.
- **Gerald Misinzo (GM)**, the advisor for the research project, helped establish the research questions and served as Julius Joseph Mwanandota’s mentor during the investigation. examined the research proposal, assisted the team conducting the study in interpreting the results, and examined the manuscript’s final revisions.
- **Augustino Chengula (AC),** the advisor for the research project, served as Julius Joseph Mwanandota’s mentor. He also helped formulate the research question, reviewed and made contributions to the study design, and reviewed and made contributions to the manuscript’s final drafts.
- **Sharadhuli Kimera (SK)** reviewed the research idea and helped with the study’s design. She mentored Julius Joseph Mwanandota throughout the research process.
- The manuscript was reviewed and edited by **Daniel Mdetele, Emmanuel Muunda, Eunice Machuka, Edward Okoth, George Omondi, and George Paul Omondi**.

## Acknowledgements

The Partnership for Skills in Applied Sciences, Engineering and Technology (PASET) provided funding for this study through the Regional Scholarship and Innovation Fund (RSIF), which was given to JJM so that he could complete his doctoral studies at the Sokoine University of Agriculture, Morogoro, Tanzania, and the SACIDS Foundation for One Health’s SACIDS Africa Centre of Excellence for Infectious Diseases. The decision to publish, the design of the study, the gathering and analysis of data, or the writing of the report was all outside the purview of the funders. The study’s conclusions and findings belong to the authors and may not necessarily reflect the opinions of the sponsors.

## Notes

### Competing Interest Statement

The authors have declared no competing interest.

## References.

1. Eloiflin RJ, Boyer M, Kwiatek O, Guendouz S, Loire E, Almeida RS de, et al. Evolution of attenuation and risk of reversal in peste des petits ruminants vaccine strain Nigeria 75/1. Viruses. 2019 Aug 1;11(8).

2. Mantip S, Sigismeau A, Shamaki D, Woma TY, Kwiatek O, Libeau G, et al. Molecular epidemiology of peste des petits ruminants virus in Nigeria: An update. Transbound Emerg Dis. 2021 May 6;

3. Bao J, Wang Q, Li L, Liu C, Zhang Z, Li J, et al. Evolutionary dynamics of recent peste des petits ruminants virus epidemic in China during 2013–2014. Virology [Internet]. 2017 Oct;510:156–64. Available from: https://linkinghub.elsevier.com/retrieve/pii/S0042682217302362

4. Ahaduzzaman M. Peste des petits ruminants (PPR) in Africa and Asia: A systematic review and meta-analysis of the prevalence in sheep and goats between 1969 and 2018. Vet Med Sci. 2020 Nov 1;6(4):813–33.

5. Tounkara K, Bataille A, Adombi CM, Maikano I, Djibo G, Settypalli TBK, et al. First genetic characterization of Peste des Petits Ruminants from Niger: On the advancing front of the Asian virus lineage. Transbound Emerg Dis. 2018 Oct 1;65(5):1145–51.

6. Dundon WG, Diallo A, Cattoli G. Peste des petits ruminants in Africa: a review of currently available molecular epidemiological data, 2020. Arch Virol [Internet]. 2020 Oct 1;165(10):2147–63. Available from: 10.1007/s00705-020-04732-1

7. Mao, L.; Li, W.; Hao F., Yang, L.; Li, J.; Sun, M.; Zhang W., Liu, M.; Luo, X.; Cheng Z. Research Progress on Emerging Viral Pathogens of Small Ruminants in China during the Last Decade. Viruses. 2022;14:1288.

8. Herzog CM, de Glanville WA, Willett BJ, Kibona TJ, Cattadori IM, Kapur V, et al. Pastoral production is associated with increased peste des petits ruminants seroprevalence in northern Tanzania across sheep, goats and cattle. Epidemiol Infect. 2019;147.

9. Jemberu WT, Knight-Jones TJD, Gebru A, Mekonnen SA, Yirga A, Sibhatu D, et al. Economic impact of a peste des petits ruminants outbreak and vaccination cost in northwest Ethiopia. Transbound Emerg Dis. 2022;

10. FAO, OIE. Global strategy for the control and eradication of PPR. International conference for the control and eradication of Peste des Petits Ruminants (PPR). Abidjan, Ivory Coast. [Internet]. 2015. Available from: https://www.woah.org/app/uploads/2021/12/ppr-global-strategy-avecannexes-2015-03-28.pdf

11. Nkamwesiga J, Korennoy F, Lumu P, Nsamba P, Mwiine FN, Roesel K, et al. Spatio-temporal cluster analysis and transmission drivers for Peste des Petits Ruminants in Uganda. Transbound Emerg Dis. 2022;

12. Motta P, Porphyre T, Handel I, Hamman SM, Ngu Ngwa V, Tanya V, et al. Implications of the cattle trade network in Cameroon for regional disease prevention and control. Sci Rep. 2017;7(August 2016):1–13.

13. Krieger N. Place, Space, and Health: GIS and Epidemiology. 2003;384–5.

14. Kao RR, Danon L, Green DM, Kiss IZ. Demographic structure and pathogen dynamics on the network of livestock movements in Great Britain. Proc R Soc B Biol Sci. 2006 Aug 22;273(1597):1999–2007.

15. Bataille A, Salami H, Seck I, Lo MM, Ba A, Diop M, et al. Combining viral genetic and animal mobility network data to unravel peste des petits ruminants transmission dynamics in West Africa. Fournié G, editor. PLOS Pathog [Internet]. 2021 Mar 18;17(3):e1009397. Available from: https://dx.plos.org/10.1371/journal.ppat.1009397

16. Rony MS, Rahman AKMA, Alam MM, Dhand N, Ward MP. Peste des Petits Ruminants risk factors and space–time clusters in Mymensingh, Bangladesh. Transbound Emerg Dis. 2017;64(6):2042–8.

17. Carpenter TE. Methods to investigate spatial and temporal clustering in veterinary epidemiology. Prev Vet Med [Internet]. 2001;48(4). Available from: 10.1016/S0167-5877(00)00199-9

18. Kanankege KST. An Introductory Framework for Choosing Spatiotemporal Analytical Tools in Population-Level. 2020;7(July):1–16.

19. González Gordon, L., Porphyre T, Muhanguzi, D., Muwonge, A., Boden, L., & de C Bronsvoort B. A scoping review of foot-and-mouth disease risk, based on spatial and spatio-temporal analysis of outbreaks in endemic settings. Transbound Emerg Dis. 2022;1–18.

20. Kraemer MUG, Reiner RC, Bhatt S. Causal Inference in Spatial Mapping. Trends Parasitol [Internet]. 2019;35(10):743–6. Available from: 10.1016/j.pt.2019.06.005

21. Perez L, Dragicevic S. An agent-based approach for modeling dynamics of contagious disease spread. Int J Health Geogr. 2009;8(1):1–17.

22. Idoga ES, Armson B, Alafiatayo R, Ogwuche A, Mijten E, Ekiri AB, et al. A Review of the Current Status of Peste des Petits Ruminants Epidemiology in Small Ruminants in Tanzania. Front Vet Sci. 2020;7(November):1–10.

23. Mdetele DP, Komba E, Seth MD, Misinzo G, Kock R, Jones BA. Review of Peste des Petits Ruminants Occurrence and Spread in Tanzania. Animals [Internet]. 2021 Jun 7;11(6):1698. Available from: https://www.mdpi.com/2076-2615/11/6/1698

24. Yizengaw L, Molla W, Temesgen W. Sero-Prevalence and Risk Factor of. bioRxiv. 2021.

25. Sowjanya Kumari S, Bhavya AP, Akshata N, Kumar K V., Bokade PP, Suresh KP, et al. Peste Des Petits Ruminants in Atypical Hosts and Wildlife: Systematic Review and Meta-Analysis of the Prevalence between 2001 and 2021. Arch Razi Inst. 2021;76(6):1589–606.

26. Kinimi E, Odongo S, Muyldermans S, Kock R, Misinzo G. Paradigm shift in the diagnosis of peste des petits ruminants: Scoping review. Acta Vet Scand [Internet]. 2020;62(1):1–14. Available from: 10.1186/s13028-020-0505-x

27. Dou Y, Liang Z, Prajapati M, Zhang R, Li Y, Zhang Z. Expanding Diversity of Susceptible Hosts in Peste Des Petits Ruminants Virus Infection and Its Potential Mechanism Beyond. Front Vet Sci. 2020 Feb 28;7(February):1–13.

28. Tricco AC, Lillie E, Zarin W, O’Brien KK, Colquhoun H, Levac D, et al. PRISMA extension for scoping reviews (PRISMA-ScR): Checklist and explanation. Vol. 169, Annals of Internal Medicine. American College of Physicians; 2018. p. 467–73.

29. Arksey H, O’Malley L. Scoping studies: Towards a methodological framework. Int J Soc Res Methodol Theory Pract. 2005 Feb;8(1):19–32.

30. Delegates WA of O. RESOLUTIONS Adopted by the World Assembly of OIE Delegates During the 89th General Session 23 – 26 May 2022. 2022;(May).

31. Bouchemla F, Agoltsov VA, Popova OM, Padilo LP. Assessment of the peste des petits ruminants world epizootic situation and estimate its spreading to Russia. Vet World. 2018;11(5):612–9.

32. Fritz CE, Schuurman N, Robertson C, Lear S. A scoping review of spatial cluster analysis techniques for point-event data. 2013;7(2):183–98.

33. R Core Team. R: A language and environment for statistical computing [Computer software] [Internet]. VIENA: R Foundation for Statistical Computing; 2024. Available from: https://www.r-project.org/

34. Draw T, Maps G. Package ‘ maps ’. 2022;

35. Package T. Package ‘ dplyr ’. 2023;

36. Create T, Data E, Using V, Description G. Package ‘ggplot2’. 2022;

37. Balamurugan V, Kumar KV, Dheeraj R, Kurli R, Suresh KP, Govindaraj G, et al. Temporal and spatial epidemiological analysis of peste des petits ruminants outbreaks from the past 25 years in sheep and goats and its control in india. Viruses MDPI AG; Mar 1, 2021.

38. Odhiambo JN, Dolan CB, Troup L, Rojas NP. Spatial and spatio-Temporal epidemiological approaches to inform COVID-19 surveillance and control: A systematic review of statistical and modelling methods in Africa. BMJ Open. 2023;13(1):1–8.

39. Ma J, Jianhua X, Han L, Xiang G, Hao C, Hongbin W. Spatiotemporal pattern of peste des petits ruminants and its relationship with meteorological factors in China. Prev Vet Med. 2017;147(May):194–8.

40. Niu B, Liang R, Zhang S, Sun X, Li F, Qiu S, et al. Spatiotemporal characteristics analysis and potential distribution prediction of peste des petits ruminants (PPR) in China from 2007–2018. Transbound Emerg Dis [Internet]. 2022 Sep 3;69(5):2747–63. Available from: 10.1111/tbed.14426

41. Carpenter TE. Methods to investigate spatial and temporal clustering in veterinary epidemiology. Prev Vet Med [Internet]. 2001 Mar;48(4):303–20. Available from: https://linkinghub.elsevier.com/retrieve/pii/S0167587700001999

42. Mergenthaler C, van Gurp M, Rood E, Bakker M. The study of spatial autocorrelation for infectious disease epidemiology decision-making: a systematized literature review. CAB Rev Perspect Agric Vet Sci Nutr Nat Resour. 2022;2022(2022).

43. Zeng Z, Gao S, Wang HN, Huang LY, Wang XL. A predictive analysis on the risk of peste des petits ruminants in livestock in the Trans-Himalayan region and validation of its transboundary transmission paths. PLoS One [Internet]. 2021;16(9 September):1–17. Available from: 10.1371/journal.pone.0257094

44. Niu B, Liang R, Zhou G, Zhang Q, Su Q, Qu X, et al. Prediction for Global Peste des Petits Ruminants Outbreaks Based on a Combination of Random Forest Algorithms and Meteorological Data. Front Vet Sci [Internet]. 2021 Jan 7;7(January):1–14. Available from: https://www.frontiersin.org/articles/10.3389/fvets.2020.570829/full

45. Abdrakhmanov SK, Mukhanbetkaliyev YY, Sultanov AA, Yessembekova GN, Borovikov SN, Namet A, et al. Mapping the risks of the spread of peste des petits ruminants in the Republic of Kazakhstan. Transbound Emerg Dis. 2021;(July):1–10.

46. Rahman AKMA, Islam SS, Sufian MA, Talukder MH, Ward MP, Martínez-López B. Peste des Petits Ruminants Risk Factors and Space-Time Clusters in Bangladesh. Front Vet Sci. 2021 Jan 25;7.

47. Zhang S, Li N, Xu M, Huang ZYX, Gu Z, Yin S. Urbanization and Habitat Characteristics Associated with the Occurrence of Peste des Petits Ruminants in Africa. Sustain. 2022;14(15):1–11.

48. Folitse E, Amemor E, Nyarku Rejoice E, Emikpe B, Tasiame W. Pattern of peste des petits ruminants (PPR) distribution in Ghana (2005–2013). Bulg J Vet Med [Internet]. 2017;20(1):51–7. Available from: http://tru.uni-sz.bg/bjvm/BJVM-March_2017_p.51-57.pdf

49. Agoltsov VA, Podshibyakin DV, Padilo LP. Analysis of peste des petits ruminants virus spread and the risk of its introduction into the territory of the Russian Federation. 2022;15:1610–6.

50. Bouchemla F, Sherasiya A, Benseghir H, MimouniA. World epidemiological meta-analyses of peste des petits ruminants (1997-2017). Rev Sci Tech l’OIE [Internet]. 2020;39(3):871–81. Available from: https://doc.oie.int/dyn/portal/index.xhtml?page=alo&aloId=42247

51. Herzog CM, Glanville WA De, Willett BJ, Cattadori IM, Kapur V, Hudson PJ, et al. Peste des petits ruminants Virus Transmission Scaling and Husbandry Practices That Contribute to among Sheep, Goats, and Cattle in Northern Tanzania. Viruses [Internet]. 2020;12(930):20. Available from: www.mdpi.com/journal/viruses

52. Muniraju M, Munir M, Parthiban AR, Banyard AC, Bao J, Wang Z, et al. Molecular Evolution of Peste des Petits Ruminants Virus. Emerg Infect Dis [Internet]. 2014 Dec;20(12):2023–33. Available from: http://www.nc.cdc.gov/eid/article/20/12/14-0684_article.htm

53. Fournié G, Waret-Szkuta A, Camacho A, Yigezu LM, Pfeiffer DU, Roger F. A dynamic model of transmission and elimination of peste des petits ruminants in Ethiopia. Proc Natl Acad Sci U S A [Internet]. 2018 Aug 14;115(33):8454–9. Available from: https://pnas.org/doi/full/10.1073/pnas.1711646115

54. Yessenbayev K, Mukhanbetkaliyev Y, Yessembekova G, Kadyrov A, Sultanov A, Bainiyazov A, et al. Simulating the Spread of Peste des Petits Ruminants in Kazakhstan Using the North American Animal Disease Spread Model. Diaz D, editor. Transbound Emerg Dis [Internet]. 2023;2023:7052175. Available from: 10.1155/2023/7052175

55. Fathelrahman EM, Reeves A, Mohamed MS, Ali YME, Awad AIE, Bensalah OK, et al. Epidemiology and cost of peste des petits ruminants (Ppr) eradication in small ruminants in the united arab emirates—disease spread and control strategies simulations. Animals. 2021;11(9).

56. Zhang S, Liang R, Yang Q, Yang Y, Qiu S, Zhang H, et al. Epidemiologic and import risk analysis of Peste des petits ruminants between 2010 and 2018 in India. BMC Vet Res [Internet]. 2022;18(1):1–15. Available from: 10.1186/s12917-022-03507-x

57. Gao X, Liu T, Zheng K, Xiao J, Wang H. Spatio-temporal analysis of peste des petits ruminants outbreaks in PR China (2013–2018): Updates based on the newest data. Transbound Emerg Dis. 2019;66(5):2163–70.

58. Assefa A, Tibebu A, Bihon A, Yimana M. Global ecological niche modelling of current and future distribution of peste des petits ruminants virus (PPRv) with an ensemble modelling algorithm. Transbound Emerg Dis. 2021;68(6):3601–10.

59. Michael D. Mitchell, Walter E. Beyeler, Patrick Finley, Melissa Finley DVM P. Modeling Peste des Petits Ruminants (PPR) Disease Propagation and Control Strategies Using Memoryless State. 2017;1(2). Available from: 10.22158/asir.v1n2p90

60. Henokmulatu Metaferia. Spatio-Temporal Distribution of Peste Des Petitis Ruminants Disease Outbreaks from 2006 to 2020 in Sheep and Goats at West Hararghe Zone, Eastern Ethiopia [Internet]. HARAMAYA UNIVERSITY, HARAMAYA; 2022. Available from: http://ir.haramaya.edu.et/hru/bitstream/handle/123456789/5555/AfterDefense2(1).pdf?sequence=1

61. ElArbi AS, Kane Y, Metras R, Hammami P, Ciss M, Beye A, et al. PPR Control in a Sahelian Setting: What Vaccination Strategy for Mauritania? Front Vet Sci. 2019;6(July):1–18.

62. Ruget AS, Tran A, Waret-Szkuta A, Moutroifi YO, Charafouddine O, Cardinale E, et al. Spatial Multicriteria Evaluation for Mapping the Risk of Occurrence of Peste des Petits Ruminants in Eastern Africa and the Union of the Comoros. Front Vet Sci. 2019 Dec 12;6.

63. Benfield CTO, Hill S, Shatar M, Shiilegdamba E, Damdinjav B, Fine A, et al. Molecular epidemiology of peste des petits ruminants virus emergence in critically endangered Mongolian saiga antelope and other wild ungulates. Virus Evol. 2021;7(2).

64. Hammami P, Lancelot R, Domenech J, Lesnoff M. Ex-ante assessment of different vaccination-based control schedules against the peste des petits ruminants virus in sub-Saharan Africa. PLoS One. 2018;13(1):1–20.

65. Haroun M, Abdulla NM, Habeb M. Peste Des Petits Ruminants: A First Retrospective Investigation Among Susceptible Animal Species in Qatar. 2021;1–19. Available from: 10.21203/rs.3.rs-371540/v1

66. Rahman AU, Dhama K, Ali Q, Hussain I, Oneeb M, Chaudhary U, et al. Peste des petits ruminants in large ruminants, camels and unusual hosts. Vet Q [Internet]. 2020;40(1):35–42. Available from: 10.1080/01652176.2020.1714096

67. Cao Z, Jin Y, Shen T, Xu F, Li Y. Risk factors and distribution for peste des petits ruminants (PPR) in Mainland China. Small Rumin Res. 2018 May 1;162:12–6.

68. Vashist VS, Yadav AK, Rajak KK, Sekar SC, Ramakrishnan MA, Muthuchelvan D, et al. Flock Level Economic Loss Due to Peste Des Petits Ruminants Outbreak in Transhumance Sheep and Goat Population in Himachal Pradesh (India). Indian J Anim Res. 2021;I(Of):1–7.

69. Aguilar XF, Mahapatra M, Begovoeva M, Kalema-Zikusoka G, Driciru M, Ayebazibwe C, et al. Peste des petits ruminants at the wildlife–livestock interface in the northern Albertine Rift and Nile Basin, East Africa. Viruses. 2020;12(3).

70. Pruvot M, Fine AE, Hollinger C, Strindberg S, Damdinjav B, Buuveibaatar B, et al. Outbreak of peste des petits ruminants virus among critically endangered mongolian saiga and other wild ungulates, Mongolia, 2016-2017. Emerg Infect Dis. 2020;26(1):51–62.

71. Gao S, Xu GY, Zeng Z, Lv JN, Huang LY, Wang HN, et al. Transboundary spread of peste des petits ruminants virus in western China: A prediction model. PLoS One [Internet]. 2021;16(9 September):1–14. Available from: 10.1371/journal.pone.0257898

72. Hammami P, Lancelot R, Lesnoff M. Modelling the dynamics of post-vaccination immunity rate in a population of sahelian sheep after a vaccination campaign against peste des petits ruminants virus. PLoS One. 2016;11(9):1–24.

73. Polonsky JA, Baidjoe A, Kamvar ZN, Cori A, Durski K, John Edmunds W, et al. Outbreak analytics: A developing data science for informing the response to emerging pathogens. Philos Trans R Soc B Biol Sci. 2019;374(1776).

74. Bouayad S, de Bellefon M-P. Spatial autocorrelation indices. Handb Spat Anal Theory Appl with R. 2018;52–70.

75. Pascual M, Dobson A. Seasonal patterns of infectious diseases. PLoS Med. 2005;2(1):0018–20.

76. Asil RM, Ludlow M, Ballal A, Alsarraj S, Ali WH, Mohamed BA, et al. First detection and genetic characterization of peste des petits ruminants virus from dorcas gazelles “Gazella dorcas” in the Sudan, 2016-2017. Arch Virol [Internet]. 2019;164(10):2537–43. Available from: 10.1007/s00705-019-04330-w

77. Jones BA, Mahapatra M, Mdetele D, Keyyu J, Gakuya F, Eblate E, et al. Peste des petits ruminants virus infection at the wildlife–livestock interface in the greater serengeti ecosystem, 2015–2019. Viruses. 2021 May 1;13(5).

78. Amprako L, Karg H, Roessler R, Provost J, Akoto-Danso EK, Sidibe S, et al. Vehicular livestock mobility in West Africa: Seasonal traffic flows of cattle, sheep, and goats across bamako. Sustain. 2021;13(1):1–16.

79. Omondi I, Galiè A, Teufel N, Loriba A, Kariuki E, Baltenweck I. Women’s Empowerment and Livestock Vaccination: Evidence from Peste des Petits Ruminants Vaccination Interventions in Northern Ghana. Animals. 2022;12(6).

80. Wahome R. www.roavs.com Analysis of small ruminants’ pastoral management practices as risk factors of Peste des petits ruminants (PPR) spread in Turkana District, Kenya. 2013;

81. Spiegel KA, Havas KA. The socioeconomic factors surrounding the initial emergence of peste des petits ruminants in Kenya, Uganda, and Tanzania from 2006 through 2008. Transbound Emerg Dis. 2019;66(2):627–33.

82. Niedbalski W, Fitzner A, Bulenger K, Kȩsy A. Peste des petits ruminants - crucial challenges for the successful disease eradication. Med Weter. 2022;78(4):159–64.

83. Real LA, Biek R. Infectious Disease Modeling and the Dynamics. Wildl Emerg Zoonotic Dis Biol Circumstances, Consequences Cross-Species Transm. 2007;33–49.

84. Nkamwesiga J, Coffin-Schmitt J, Ochwo S, Mwiine FN, Palopoli A, Ndekezi C, et al. Identification of peste des petits ruminants transmission hotspots in the Karamoja subregion of Uganda for targeting of eradication interventions. Front Vet Sci. 2019 Jul 5;6(JUL).

85. Fine AE, Pruvot M, Benfield CTO, Caron A, Cattoli G, Chardonnet P, et al. Eradication of Peste des Petits Ruminants Virus and the Wildlife-Livestock Interface. Front Vet Sci. 2020;7(March):1–8.

86. Chowell G, Rothenberg R. Spatial infectious disease epidemiology: On the cusp. BMC Med. 2018;16(1):1–5.

87. Folitse E, Amemor E, Nyarku Rejoice E, Emikpe B, Tasiame W. Pattern of peste des petits ruminants (PPR) distribution in Ghana (2005–2013). Bulg J Vet Med [Internet]. 2017;20(1):51–7. Available from: http://tru.uni-sz.bg/bjvm/BJVM-March_2017_p.51-57.pdf

88. Bouchemla F, Sherasiya A, Benseghir H, Mimouni A. World epidemiological meta-analyses of peste des petits ruminants (1997-2017). Rev Sci Tech l’OIE [Internet]. 2020 Dec 1;39(3):871–81. Available from: https://doc.oie.int/dyn/portal/index.xhtml?page=alo&aloId=42247

89. Kivaria FM, Kwiatek O, Kapaga AM, Swai ES, Libeau G, Moshy W, et al. The incursion, persistence and spread of peste des petits ruminants in Tanzania: Epidemiological patterns and predictions. Onderstepoort J Vet Res [Internet]. 2014;81(1). Available from: https://ojvr.org/index.php/ojvr/article/view/593/914

